# Goat Discrimination of Emotional Valence in the Human Voice

**DOI:** 10.1101/2023.05.26.542400

**Authors:** Marianne A. Mason, Stuart Semple, Harry H. Marshall, Alan G. McElligott

## Abstract

Reading another animal’s emotional state can enable receivers to anticipate their behavioural motivations, which is important in guiding interactions with that individual. For species living closely alongside humans, the emotional cues that people express can be almost as informative as those of conspecifics. Goats can discriminate differences in emotional valence present in another goat’s calls, and we investigated whether this ability extends to human speech. We presented goats with a habituation-dishabituation-rehabituation paradigm, where they experienced multiple playbacks of a familiar or unfamiliar human voice conveying a single emotional valence (e.g., angry; habituation phase), before the valence of the voice changed (e.g., happy; dishabituation phase) and then reversed again in-line with the habituation phase (e.g., angry; rehabituation phase). Over the habituation phase, goat behavioural responses decreased, showing evidence of having habituated to the playback stimuli presented. Following a change in emotional valence (dishabituation phase), although goats were overall less likely to respond, those that did looked for longer, suggesting they had perceived the shift in emotional content conveyed in human voice playbacks. We found no changes in physiological arousal (heart rate or heart rate variability) with shifts in playback valence. Goats, as a domesticated species, may have developed a sensitivity to our cues over their long association with humans, but the differences in individual behaviour towards the playback paradigm could highlight a role for learning and individual experience in particular on interspecific emotional communication.

## 1. INTRODUCTION

Emotional processes help tailor an animal’s behaviour adaptively towards its environment (Kremer et al., 2020), for example, motivating movement towards fitness-enhancing stimuli, such as mates and foraging resources, and away from fitness-threatening ones, like predators. Such responses incorporate neural, physiological, behavioural and (sometimes) subjective components (supposing a conscious feeling of emotions experienced; Špinka, 2012; Kremer et al., 2020). The behavioural component is often exaggerated and somewhat distinct, leading many researchers to conclude that while emotional displays generally serve a direct function (e.g., moving away from danger), in a social species, they may play additional roles in communication (Shariff & Tracy, 2011). Indeed, receivers may benefit from reading another individual’s emotional state as it enables them to predict a signaller’s future behaviour, important in guiding interactions with that individual (Schmidt & Cohn, 2001).

One common way of understanding emotion is to split these responses across two or more dimensions (known as dimensional approaches), usually valence (positive or negative, e.g., happiness to sadness) and arousal (the intensity of the emotion experienced, e.g., calm to excited; Mendl et al., 2010). Although vocalisations are generally inherently communicative in nature, emotional experiences differing in both these dimensions create measurable differences in animal vocalisations, with calls in more arousing situations tending to be louder, of higher frequency and produced at a greater rate (Briefer, 2012, 2020). Conversely, shifts in vocal features relative to valence appear more species-specific, with different types of call being favoured in different social and emotional contexts (Briefer, 2012, 2020). For example, laughter and crying in humans tend to be linked to positive and negative emotions respectively. However, it has been suggested positive vocalisations tend to be shorter than negative ones, with a lower and less variable fundamental frequency (relates to pitch; Briefer, 2020).

Vocalisations are considered to match emotional state more closely in non-human animals than in humans, with this relationship becoming more complex in our own species owing to the greater conscious control we have over our vocal apparatus (Jürgens, 2009; Briefer, 2012). However, the link between emotional state and vocal communication has not been entirely lost, with emotional cues remaining clearly evident in our non-verbal utterances (e.g., crying and laughter) and speech (Ackermann et al., 2014; Bryant, 2018, 2021). Similarities in how emotion is conveyed vocally across taxonomic groups may to an extent enable animals to apply rules from their own species-specific emotional communication systems to discriminate analogous heterospecific cues (Faragó et al., 2014; Filippi et al., 2017; Bryant, 2021). This accuracy in which an animal can discriminate another species emotional cues may be further developed through experience with the species in question (Kitchen et al., 2010; Merola et al., 2014; Scheumann et al., 2014; Barber et al., 2016).

For species living alongside humans, the cues we express not only advertise the level of threat we may pose, but in some cases, the potential benefits that our presence or actions may offer (e.g., feeding opportunities; Goumas et al., 2022; Cram et al., 2022). The most appropriate response an animal can give towards a given person (e.g., ignore, approach or avoid) will vary among people and over time, based on a particular person’s current motivations. Accordingly, a number of species appear to have developed a sensitivity to broad demographic cues, such as a person’s ethnicity, gender and age group (Bates et al., 2007; McComb et al., 2014; McIvor et al., 2022), as well as towards more proximate cues, such as particular vocal signals (e.g., whistling), gaze direction (Carter et al., 2008; Spottiswoode et al., 2016; Goumas et al., 2019; Cram et al., 2022) and, moreover, emotional expressions. Our emotional cues offer clues as to how we may behave in the immediate future, which if perceived, can be used to anticipate positive and negative events as a result of interacting with certain people (Schmidt & Cohn, 2001). If sensitive to these cues and for domesticated species particularly, which have relied on us for their care over thousands of generations (MacHugh et al., 2017) our emotional expressions may in turn affect animal emotional experiences and welfare (Merkies et al., 2013; Proops et al., 2018; Smith et al., 2018; Lange et al., 2020). Companion animals have been shown to perceive human emotional cues conveyed in multiple sensory channels, including in combination (using visual and vocal cues: Albuquerque et al., 2016; Nakamura et al., 2018; Quaranta et al., 2020). However, far less is known about these abilities in livestock.

Unlike species domesticated for companionship or work, livestock have been bred largely for their products (e.g., meat, milk and hair), so may be expected to be under weaker selection to discriminate between and respond to our cues, including emotional ones (MacHugh et al., 2017; Jardat & Lansade, 2021). Despite this, some livestock have been shown to read a variety of human cues sometimes showing comparable levels of performance to companion animals. For example, pigs (*Sus scrofa domesticus*) respond differently to human voices based on their emotional valence (Maigrot et al., 2022) and cows (*Bos taurus*) can discriminate chemical cues, preferring to interact with sweat odours collected from humans in neutral, but not in stressful situations (Destrez et al., 2021).

Goats (*Capra hircus*), a species primarily domesticated for their products (MacHugh & Bradley, 2001; but see pack goats, Sutliff, 2019) respond to a variety of human cues. For example, when confronted with an unsolvable task, goats altered their human-directed gazing behaviour in relation to the perceived level of attention they received (Nawroth et al., 2016a). This not only indicates that can goats discriminate our attentional cues, but these gaze alternation behaviours have been interpreted as attempts at human-directed communication. Goats can also use our gestural cues to solve tasks (pointing/ tapping: Kaminski et al., 2005; Nawroth et al., 2015, 2020) and even learn socially from humans (Nawroth et al., 2016b). Moreover, they are sensitive to emotional cues, discriminating between human, as well as conspecific positive and negative facial expressions (Bellegarde et al., 2017; Nawroth et al., 2018). That being said, the ability of goats and other ungulate livestock themselves to produce complex, emotive facial expressions is constrained by their relatively limited facial musculature and the morphology of these expressions appear poorly conserved across taxa (Waller & Micheletta, 2013).

Goats are a highly vocal species, with calls encoding a range of information, including the caller’s age, body size, sex, social environment, individual identity (Briefer & McElligott 2011a, 2011b, 2012) and importantly, their emotional state (Briefer et al., 2015a). Such variability provides ample substrate for making related distinctions and indeed, goats have been shown to discriminate identity information (Briefer & McElligott, 2011a; Briefer et al., 2012; Pitcher et al., 2017) and moreover, emotional valence (Baciadonna et al., 2019) conveyed in conspecific calls. To test whether they can discriminate analogous emotional cues in the human voice, we presented goats with a playback paradigm to investigate whether subjects could perceive shifts in valence, specifically, anger versus happiness conveyed in human speech.

Goats listened to a series of nine voice playbacks expressing either a positive (happy) or a negative (angry) valence (habituation phase), before the valence of playbacks was changed (e.g., from positive to negative: dishabituation phase) and then reversed in line with that of the habituation phase (based on Charlton et al., 2007, 2011; Baciadonna et al., 2019). We predicted that if goats could discriminate emotional content conveyed in the human voice, they would dishabituate, looking faster and for longer towards the source of the sound, following the first shift in valence. We also expected potential changes at physiological levels with heart rate increasing and heart rate variability (HRV) decreasing after a shift in valence, indicating an increase in arousal (Briefer et al., 2015a; Baciadonna et al., 2020). Alternatively, goats may also express differences in arousal based on the valence that the voice conveyed (Baciadonna et al., 2019). In dogs (*Canis lupus familiaris*) and horses (*Equus caballus*), familiarity with the individual conveying the emotional cues appears to facilitate their discrimination (e.g., Merola et al., 2014; Briefer et al., 2017), and thus goat responses were also investigated in relation to whether voice playbacks were taken from familiar or unfamiliar people. We expected if familiar voices are easier to discriminate, goats would respond faster and for longer when presented with familiar over unfamiliar voices following a change in valence, perhaps accompanied with increases in physiological arousal (again, indicated through cardiac measures).

## 2. METHODS

### 2.1. Study Site & Sample Population

We carried out experiments at Buttercups Sanctuary for Goats (http://www.buttercups.org.uk/) in Kent, UK (51°13’15.7”N 0°33’05.1”E) between 6^th^ August and 2^nd^ October 2019. The sanctuary was open to visitors and features a large outdoor paddock (3.5-4 acres approximately) which goats could access throughout the day, and at night they were kept either individually or in small groups (average pen size 3.5m^2^). Animals had *ad libitum* hay, grass and water available and were provided with commercial concentrate in relation to age and condition. Our final sample size for this study was 27 subjects (13 castrated males and 14 intact females) of various breeds and ages (for further information, see Appendix Table A1) that had resided at the sanctuary for over a year and were well-habituated to human handling.

### 2.2. Collection & Preparation of Auditory Stimuli

We collected human voice samples using a Sennheiser MKH 416 P48 directional microphone in combination with a Marantz PMD-661 digital recorder (sampling rate: 48kHz, with an amplitude resolution of 16 bits in WAV format). Recordings were taken from eight speakers for which English was their first language, four of whom were familiar to subjects and four unfamiliar (two male and two female speakers in each group). Those familiar to subjects had worked or volunteered in a husbandry-related capacity at the sanctuary for at least one year. Recording sessions took place in enclosed, quiet areas to minimise background noise, although a common location in which all recordings could be collected was not possible. While recording, we held the microphone close to the speaker’s face (approximately 20cm away) and they were asked to say the phrase “hey, look over here” multiple times, first using a happy voice (positive valence) and then an angry voice (negative valence). The same phrase was used so responses would depend on the emotional content of the voice sample rather than the specific words used (Schamberg et al., 2018).

Recording quality and expression of emotional valence were evaluated for each voice sample by a single listener. If recordings were not deemed to be of sufficient quality, the recording session was repeated. Only voice samples taken from a single recording session were used in playbacks to ensure auditory environment in which samples were recorded was consistent across the entire playback. We used the three clearest samples judged to be most strongly expressing the desired emotional valence for both positive and negatively-valenced samples to construct playbacks (giving six voice samples per speaker in total). Voice samples had a mean length (± SD) of 1.43s ± 0.373, with length not differing significantly according to the emotional valence conveyed (linear mixed model: β ± s.e. = 0.102 ± 0.085, t_(39)_ = 1.20, *p* = 0.238).

We assembled playback sequences using Praat v.6.1. (Boersma & Weenink, 2019), with these comprising a series of voice samples from a single speaker, with each of the 13 samples followed by 20s of silence, in which time (hereafter known as the response period), behavioural and cardiac responses were measured. Mean amplitude of recordings was scaled digitally to 70dB, but when measured under field conditions, there was a very small absolute (1.23dB), but statistically significant difference in average maximum amplitude between positively- and negatively-valenced voice samples (positive: 74.18dB ± 3.026; negative: 75.41 dB ± 3.289; LMM: β ± s.e.= 1.137 ± 0.456, t = 2.49, *p* = 0.014; measured at a one meter distance using a CEMTM DT-8851 sound-level meter). Overall, playback sequences had a mean length (± SD) of 4 minutes 38s ± 4s.

### 2.3. Playback Procedure

To investigate goat ability to discriminate emotional cues in the human voice, we used a habituation-dishabituation-rehabilitation paradigm, similar to that of Baciadonna et al. (2019), who investigated their discrimination of valence in conspecific calls. In each phase, voice recordings conveyed a single emotional valence, the valence being the same in the habituation and rehabituation phases, but different in the dishabituation phase (so e.g., if valence was positive in the habituation and rehabituation phases, it was negative in the dishabituation phase). In the habituation phase three different recordings were repeated three times and combined in a random order (so nine recordings in total; H1-H9). Through repeated exposure to auditory stimuli sharing similar properties, we expected goats to lose interest, or habituate and consequently reduce responsiveness to playbacks over the course of the habituation phase. Furthermore, as different voice samples from the same speaker were used, goats were anticipated to habituate to the valence content of the samples, rather than to individual recordings (Charlton et al., 2011). The subsequent dishabituation phase consisted of three different recordings (D10-D12) conveying the opposite valence to that of the habituation phase, with recordings again combined in a random order. If a goat perceived the shift in valence, we expected them to dishabituate, or renew responsiveness towards the playbacks presented. A randomly selected single recording from the habituation phase comprised the rehabituation phase (R13). We predicted responsiveness would again decrease during this phase if shifts in behavioural and physiological measures observed during the dishabituation phase were robust and not an artefact of, for example, a chance renewal in attention (Charlton et al., 2007, 2011).

For each of the eight people providing voice samples, two playback sequences were produced: one comprising voice samples expressing a positive valence during the habituation and rehabituation phases (i.e., Happy-Angry-Happy) and one a negative valence in these phases (i.e., Angry-Happy-Angry). This produced 16 playbacks in total, eight from familiar and eight from unfamiliar people. Each goat experienced at least two trials, one using a playback of a familiar voice and one using an unfamiliar voice. The speaker’s gender and valence expressed during the habituation phase were the same across both trials. Sanctuary staff and volunteers were asked which goats appeared especially responsive to them, and therefore likely to be more familiar with their voice. When tested, these goats were subject to playbacks derived from these staff members’ voices and the rest were assigned playback stimuli semi-randomly, while ensuring half of subjects experienced playbacks of male voices, and half from females.

### 2.4. Playback Emotional Valence Validation Experiment

To verify whether the voice samples used effectively conveyed the desired emotional valence, after carrying out the playback experiments with goats, we asked a convenience sample of ten volunteers with fluency in English (five males and five females) to score all recordings according to their emotional valence (angry or happy). Participants listened to all 48 voice samples used in our experiment, which had been combined in a random order and split into three playback sequences (of 16 recordings each). Following each voice sample, participants were given five seconds of silence in which to score its emotional valence.

Participants correctly scored an average of 46.4 ± 1.27 out of 48 voice samples. 87.5% of mistakes occurred in the first playback sequence, and so it is likely this error rate is at least partially inflated as participants adapted to the experimental protocol. For most playbacks which participants had scored incorrectly, only one or two of the ten participants had made mistakes, suggesting valence had overall been effectively conveyed. There was however, one recording where only 40% of participants correctly scored valence indicating its emotional content was ambiguous, but this voice sample only featured in four out of 42 experimental trials.

### 2.5. Experimental Enclosure

We constructed the experimental enclosure from opaque metal agricultural fencing placed within the large outdoor paddock that goats could readily access throughout the day. The enclosure was divided into three sections (Fig. 1; Appendix Fig. A1). Subjects initially entered through the preparation pen, before being led into the experimental arena where trials took place. The equipment section was separated from the experimental arena by a gate comprising horizontal metal bars. A Sony CX240E video camera and speaker were positioned in this section 2.65m away from the front of the experimental arena. The Bose Soundlink Mini Bluetooth Speaker II we used to broadcast playbacks during trials has been verified as effective in reproducing the human voice (Ben-Aderet et al., 2017). We covered the camera and speaker in camouflage netting to reduce the likelihood of subjects locating the sound source. The experimenter was positioned within this section during experimental trials, with their back turned away from the goat being tested to avoid unintentionally cuing them.

**Figure 1.**
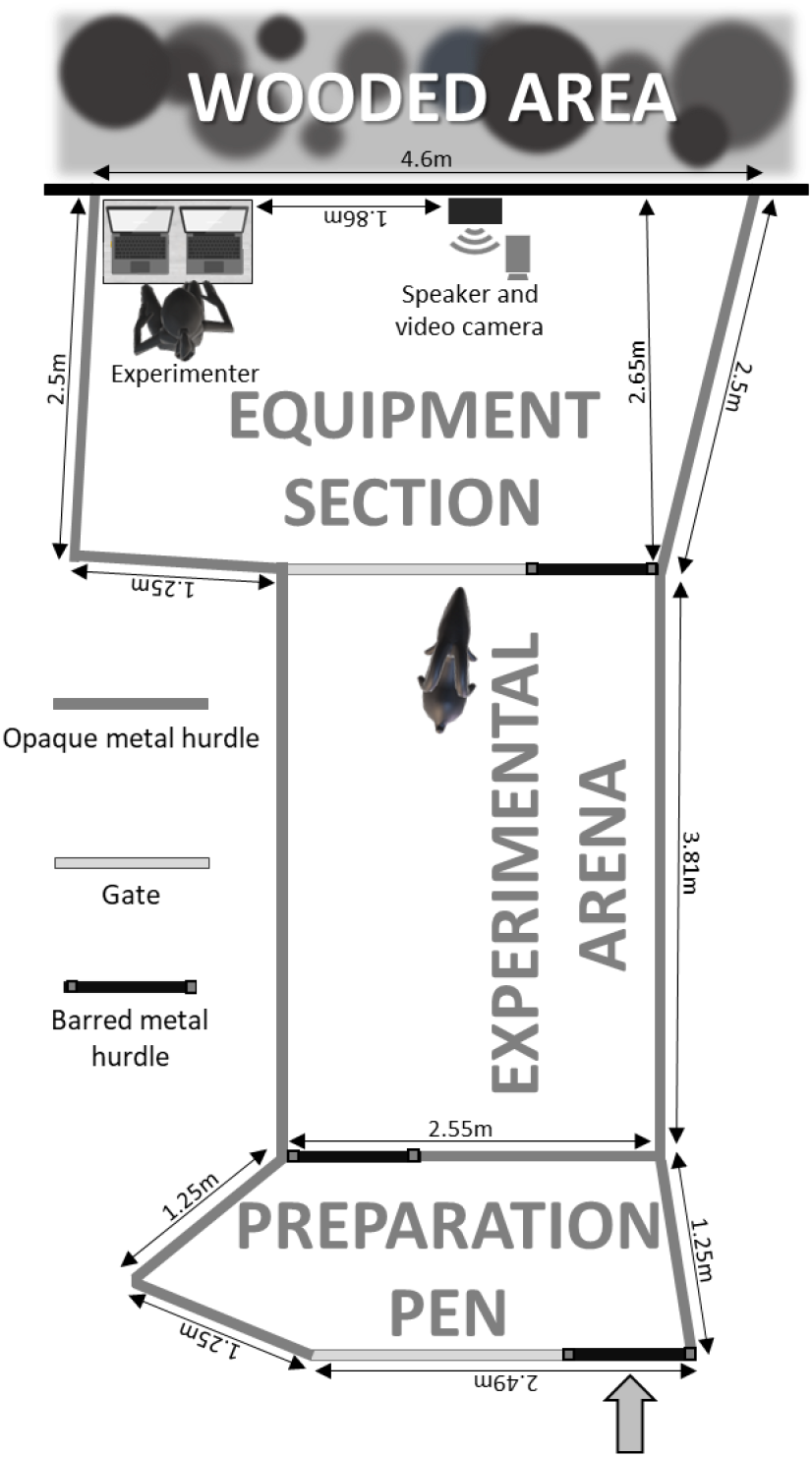
Experimental enclosure. Goats entered through the preparation pen where they were fitted with a heart rate monitor. Trials took place in the experimental arena. The speaker was positioned 2.65m away from the front of the experimental arena.

### 2.6. Ethics Statement

We conducted all animal care and experimental procedures in line with ASAB guidelines for the use of animals in research (ASAB/ ABS, 2019). Procedures were further subject to approval by the University of Roehampton’s Life Sciences Ethics Committee (Ref. LSC 19/ 280). All methods were non-invasive, and trials were a maximum of 8 minutes 5s in duration. The valence validation experiment carried out on human volunteers was approved through an amendment made to the original ethics application (Amendment 06.20).

### 2.7. Experimental Preparation & Procedure

We habituated goats to the experimental arena over two five-minute sessions taking place over consecutive days, with the final session taking place a minimum of 1 hour 50 minutes prior to the onset of experimental trials (maximum = 7 days). This variation in interval between habituation sessions and experiments was necessary as goats were not always willing to be led to the experimental enclosure or were displaced by a more dominant individual during transit. Trials themselves took place on weekdays between 11:00 am and 4:00 pm.

Subjects were first gently led to the experimental enclosure and upon entering the preparation pen, we fitted them with a Zephyr^TM^ BioHarness 3.0 affixed to a belt. During trials this device, in conjunction with AcqKnowledge v.4.4.2 software (BIOPAC System Inc.) was used to transmit live cardiac data via Bluetooth to a laptop (HP ProBook 650 G4). To improve conductance of the BioHarness to the skin, we clipped fur over the left shoulder blade (across which the BioHarness module was to be positioned) at least one week prior to onset of experiments and liberal amounts of ECG gel were applied to sensors before the belt was placed around the goat’s thorax. Similar wearable devices have been successfully implemented to monitor changes in goat heart rate and HRV in relation to a variety of factors (e.g., Briefer et al., 2015a, 2015b; Baciadonna et al., 2016, 2020), including towards conspecific emotional cues (Baciadonna et al., 2019). Once we found a clear ECG trace, the goat was led into the experimental arena.

One minute after the subject had entered the arena, we initiated data logging from the heart rate monitor and recording using the video camera. If the ECG trace had been maintained, goats were kept in the arena for a further two minutes before playbacks were initiated to habituate without human interference, but if it had been lost, the trial was paused and the BioHarness repositioned. Following initiation of the playback sequence, we marked an event in the cardiac data at the end of each recording to indicate onset of the 20s response period. This process was repeated until the end of the playback sequence, after which time we led the subject back into the preparation pen where the BioHarness was removed before they were freed. The two trials of any given subject, one of which played voice samples from a familiar, and one trial where samples were from an unfamiliar person, were separated by a minimum of seven days (mean ± SD = 9.38 days ± 3.21).

We scored videos for levels of visitor disturbance, and repeated corresponding trials when this was deemed to be too great. The repeated trial took place a minimum of 17 days after the second trial (maximum = 48 days), with 14 out of 30 goats tested having a trial repeated and four of these subjects requiring both trials to be repeated (trial number was controlled for in the statistical analysis).

### 2.8. Video Coding

We coded all experimental trials on a frame-by-frame basis using BORIS v.7.8.2. (Friard & Gamba, 2016). The majority of these were captured at a frame rate of 25 FPS, but for two trials, technical issues prevented reliable footage being taken, so behavioural data was coded from back-up footage captured by a webcam at a rate of five FPS. We specifically scored how long goats looked towards the speaker (defined as when their snout was pointed within approximately 45 degrees of the speaker), and how long it took them to look in the response periods following each of the 13 playbacks.

Data analysed was extracted by a single observer, but a second blind observer coded all trials included in the analyses to assess the reliability of the behavioural data set. This observer was not informed of experimental hypotheses, and ignorance of the playback paradigm was maintained through coding videos in absence of auditory and trial information. There was high agreement in looking duration measured by these two observers (intraclass correlation analysis; Poisson distribution assumed: R ± s.e = 0.83 ± 0.013, *p*<0.0001, n = 1000 bootstraps; R package, rptR: Stoffel et al., 2017).

### 2.9. Exclusion Criteria

Initially, we excluded one male and one female from further analysis, due to expression of abnormal behaviour and successful escape from the experimental arena, respectively. Based on criteria employed by Baciadonna et al. (2019), further trials were also excluded if they satisfied the following: 1) goats failed to respond to the first recording (i.e., did not look towards the speaker), and/ or 2) subjects did not habituate to playbacks, which was interpreted as when the looking duration after the habituation phase’s final playback exceeded by more than two times that observed in the response period following the first playback. In addition, trials in which goats did not look at the speaker for more than 10 seconds in total over the entire playback sequence were excluded as these subjects were considered not attentive enough to playbacks for their discriminatory abilities to be effectively assessed. We excluded a total of 18 trials, which included all records collected from three subjects. Ultimately, observations from 27 goats participating in 42 trials (familiar speaker = 20; unfamiliar = 22) were included in analysis.

### 2.10. Statistical Analysis

#### 2.10.1. General Model Parameters

We fitted generalised linear mixed models (GLMMs) to the response variables: looking duration, latency to look, heart rate and HRV, all measured over in the 20s response periods following each of the 13 playbacks, using R version 4.2.3. (R Core Team, 2023). To begin with, we investigated changes in each of these behavioural and physiological responses over the habituation phase (H1-H9) to examine the likelihood of habituation having taken place. Following this, we conducted pairwise comparisons to evaluate changes in relation to shifts in valence, by comparing goat responses to the last playback of the habituation phase and the first of the dishabituation phase (H9 v. D10), as well as to the last playback of the dishabituation phase and the rehabituation phase (D12 v. R13). To avoid overfitting models, we used the Akaike’s information criterion (AIC) to identify and remove variables which proved poor predictors of looking duration (choosing the model with the lowest AIC value), e.g., goat sex and speaker gender to arrive at a single set of variables for all analytical models.

Our primary variable of interest was playback number, as we wanted to examine how responses changed over the course of the habituation phase and between the habituation and dishabituation, as well as the dishabituation and rehabituation playback phases. Given familiarity with the individual expressing the emotional cues is of known importance for discriminating valence in other species (e.g., Merola et al., 2014; Briefer et al., 2017), the effect of voice familiarity (familiar v. unfamiliar) on goat responses was also investigated. Accordingly, we explored the interaction between playback number and familiarity to explicitly test whether voice familiarity affected goat ability to discriminate human emotional cues. For models where this interaction was non-significant, we removed this term to enable interpretation of the effects of playback number and familiarity independently (as recommended by Engqvist, 2005). In addition, we investigated the effect of playback order on goat responses, with this variable denoting the order in which valence is presented over the habituation, dishabituation and rehabituation phases, i.e., angry-happy-angry or happy-angry-happy. As for cardiac responses, noise present in the ECG trace sometimes prevented heart rate and HRV being measured across the entire response period. The period over which it was possible to calculate these responses was therefore referred to as the measurement period (mean = 18.10s; range = 5.96-19.46s). As longer measurement periods provide more information, this covariate can have important implications for the accuracy of cardiac measures, so was fitted to all relevant models (Reefmann et al., 2009; Briefer et al., 2015b). Finally, we nested trial number (one or two, and sometimes three or four where trials had been repeated) within goat identity and added this as a random effect to control for repeated measurements taken from each individual both within and between trials. Playback code, a unique identifier for each playback sequence was included as a separate random effect to control for the same playbacks being used over multiple trials.

#### 2.10.2. Looking Duration

As looking duration was restricted between zero and 20s, model residuals failed to conform to a normal error structure, and only few goats looked for the entire response period (median looking duration over habituation phase = 3.30s), we considered statistical approaches more typically applied to count data. As data was both zero-inflated and overdispersed (verified using the DHARMa package: Hartig, 2021), we used AIC to test the fit of multiple error structures (Poisson, zero-inflated negative binomial; glmmTMB package: Brooks et al., 2017), finding a zero-inflated negative binomial type ll model to be most appropriate for our data. This approach produces both a conditional model, which was used to predict duration values greater than zero seconds (log link and negative binomial error structure), and a zero-inflated model which predicted the probability of a zero second observation (the goat did not look) using a logit link. We unnested trial number from goat identity and removed the random effect, playback code to resolve convergence issues from models comparing goat responses between the dishabituation and rehabituation phases. Not taking into consideration such nesting patterns among related observations and the effect of playback sequence on goat responses could have made the D12-R13 comparison less robust compared to other analyses.

#### 2.10.3. Latency to Look

Latency conformed to an approximately bimodal distribution with peak abundances at zero seconds (the goat was looking from the outset of the response period) and 20s (subjects did not look throughout the response period), so this response was analysed using a binomial GLMM (glmer function, lme4 package: Bates et al., 2015). *Post hoc* comparisons for the interaction between playback number and familiarity were carried out using the emmeans package with Tukey’s corrections to control for multiple comparisons (Lenth, 2021). We removed the random effect playback code from the habituation-dishabituation phase comparison due to issues with model convergence.

#### 2.10.4. Heart Rate & Heart Rate Variability

We measured heart rate as beats per minute, and HRV as the root mean square of successive differences between heartbeats (RMSSD) measured in the 20s response period after playbacks (following Baciadonna et al., 2019). Differences in both cardiac parameters are thought to reflect shifts in physiological arousal, and have been shown to change in relation to a variety of emotional contexts in goats (e.g., Baciadonna et al., 2016, 2020; Briefer et al., 2015a, 2015b).

Although RMSSD especially is not typically considered count data, as resulting residuals from modelling cardiac responses failed to conform to a Gaussian distribution and were restricted to positive values, Poisson error structures were adopted to model both responses (lme4 package: Bates et al., 2015). Models were formally tested for dispersion using the DHARMa package (Hartig, 2021) and we fitted all models predicting goat cardiac responses with an observation-level random effect accordingly. Upon fitting models and plotting their error structures, we noticed the presence of influential observations in the heart rate data measured over the habituation phase. Using the outliers package, we identified and removed the most extreme outlier from habituation phase analyses to avoid it having a disproportionate effect on resulting model parameters. The random effect, playback code, was removed from models investigating changes in goat HRV over the habituation phase and heart rate between the habituation and dishabituation phases to resolve convergence issues.

## 3. RESULTS

### 3.1. Looking Duration

Goats looked for an increasingly shorter time and were less likely to look towards the sound source with increasing playback number over the habituation phase (conditional model: β ± s.e. = -0.075 ± 0.016, Z = -4.76, *p* = 1.98x10^-6^; zero-inflated model: β ± s.e. = 0.136 ± 0.052, Z = 2.63, *p* = 0.009, respectively; Table 1; Fig. 2). In addition, subjects were marginally less likely to look when experiencing human voices conveying a negative over a positive valence in this phase (β ± s.e. = -0.860 ± 0.486, Z= -1.77, *p* = 0.077). By contrast, we found no significant effect of familiarity with the voice presented on how likely or how long goats looked over this phase.

**Figure 2.**
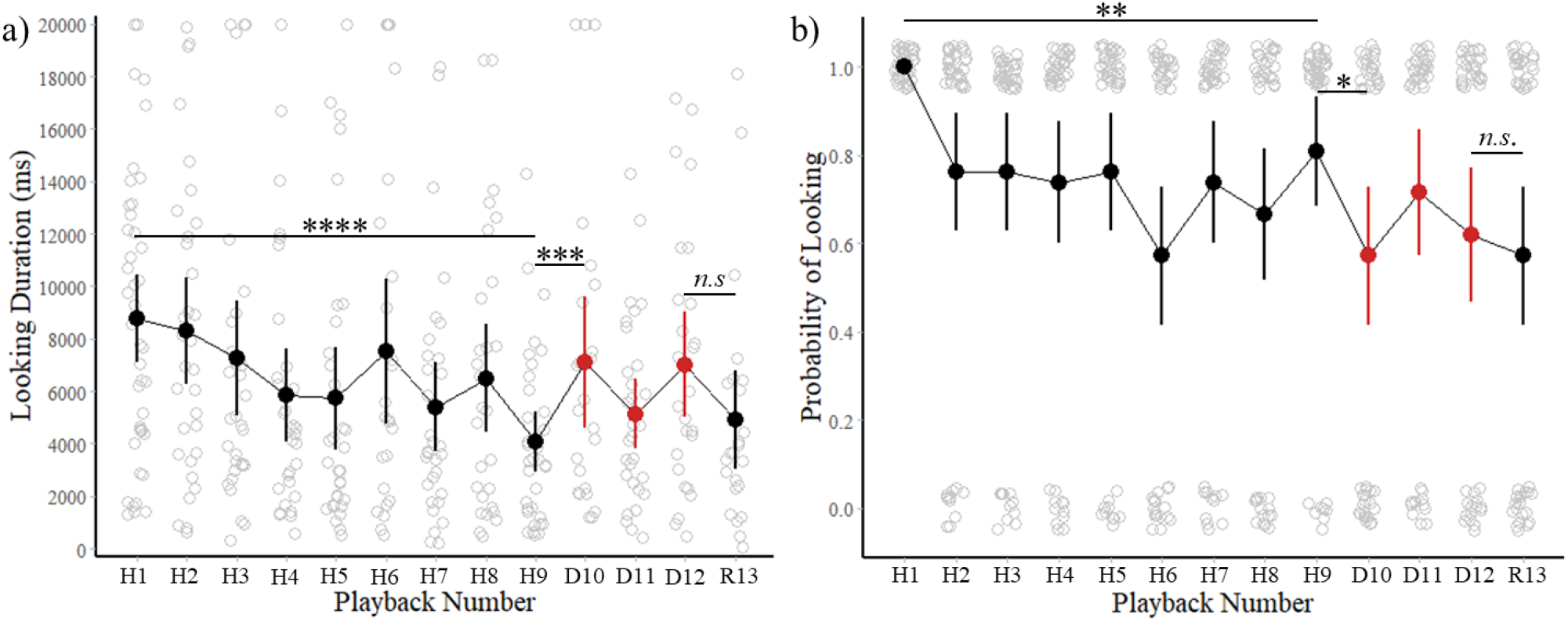
Effect of playback number on goat looking behaviours in the 20s following playbacks. Mean and 95% confidence intervals shown for a) goat looking duration values greater than zero b) the likelihood that goats looked, as a function of playback number. Goat responses in the dishabituation phase playbacks are shown in red. Comparisons are shown above and significance indicated. *n.s.* = non-significant; \**p*<0.05; \*\**p*<0.01; \*\*\**p*<0.001; \*\*\*\**p*<0.0001

**Table 1.**
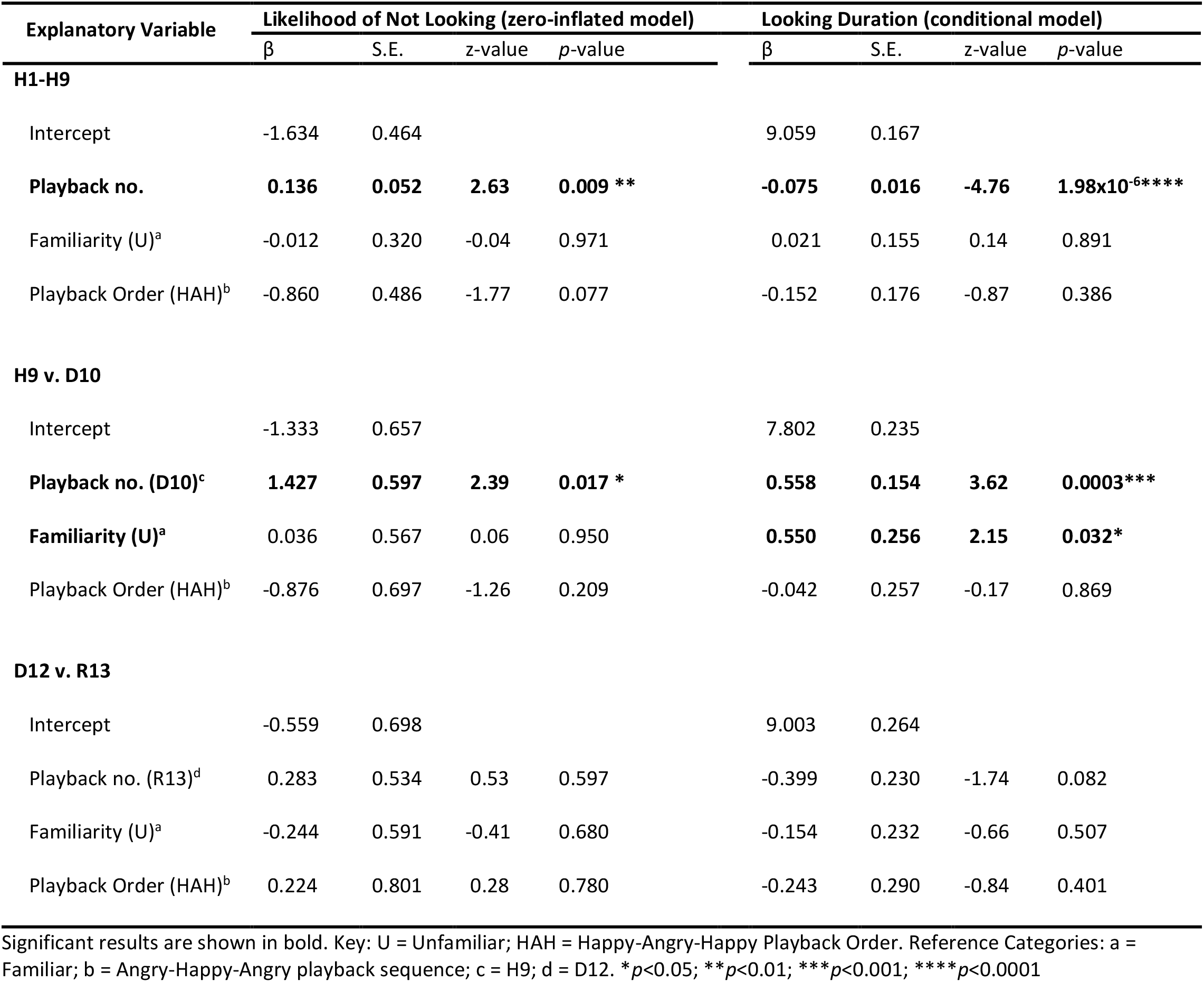
Predictors of time goats spent looking (conditional model) and likelihood that they did not look (zero-inflated model) towards the sound source following each human voice playback of the habituation phase (H1-H9), after the shift in emotional valence between the habituation and dishabituation phase (H9 v. D10) and finally, between the dishabituation and rehabituation phases (D12 v. R13). Parameter estimates come from zero-inflated type ll model.

Goats were less likely to look immediately following a change in emotional valence after the first playback of the dishabituation phase (D10), compared to the last of the habituation phase (H9; zero-inflated model: β ± s.e. = 1.427 ± 0.597, Z = 2.39, *p* = 0.017; Table 1; Fig. 2b). However, goats that looked did so for longer after this valence change (conditional model: β ± s.e. = 0.558 ± 0.154, Z = 3.62, *p* = 0.0003; Fig. 2a). When visualising the relationship between looking duration and playback number, the presence of three goats that looked for the entire 20s response period after D10 was noted (Fig. 2a). As these observations could have had a disproportionate influence on parameter estimates in the conditional model (measuring looking duration for goats that looked), we removed data from these goats but found the effect of playback number on looking duration remained significant (β ± s.e. = 0.403 ± 0.191, Z = 2.11, *p* = 0.035). Familiarity with voices presented also affected looking behaviours, with goats played unfamiliar voices looking for longer than those experiencing familiar voices over the H9 versus D10 comparison (β ± s.e. = 0.550 ± 0.256, Z = 2.15, *p* = 0.032). Between the end of the dishabituation (D12) and start of the rehabituation phases (R13), the probability that goats looked was not significantly affected, although subjects tended to look for a shorter time after R13 (β ± s.e. = -0.399 ± 0.230, Z = -1.74, *p* = 0.082; Table 1; Fig. 2). By contrast, playback order (angry-happy-angry or happy-angry-happy) had no significant effect on how long or how likely goats were to look after the dishabituation phase, and neither familiarity nor playback order had a significant effect on their responses following the rehabituation phase.

### 3.2. Latency to Look

Goats became slower to respond to playbacks as the habituation phase progressed, with time taken to look increasing at a faster rate when they were experiencing unfamiliar, compared to familiar voices (playback number x familiarity interaction: β ± s.e. = 0.059 ± 0.001, Z = 81.85, *p*<2x10^-16^****; Table 2). However, it appears that voice familiarity only had a weak (and possibly negligible) effect on goat responses, as suggested by the low effect size and lack of obvious trend when visualising this relationship (Fig. 3). The valence that the voice conveyed had no significant effect on how long it took for goats to respond to playbacks over the habituation phase.

**Figure 3.**
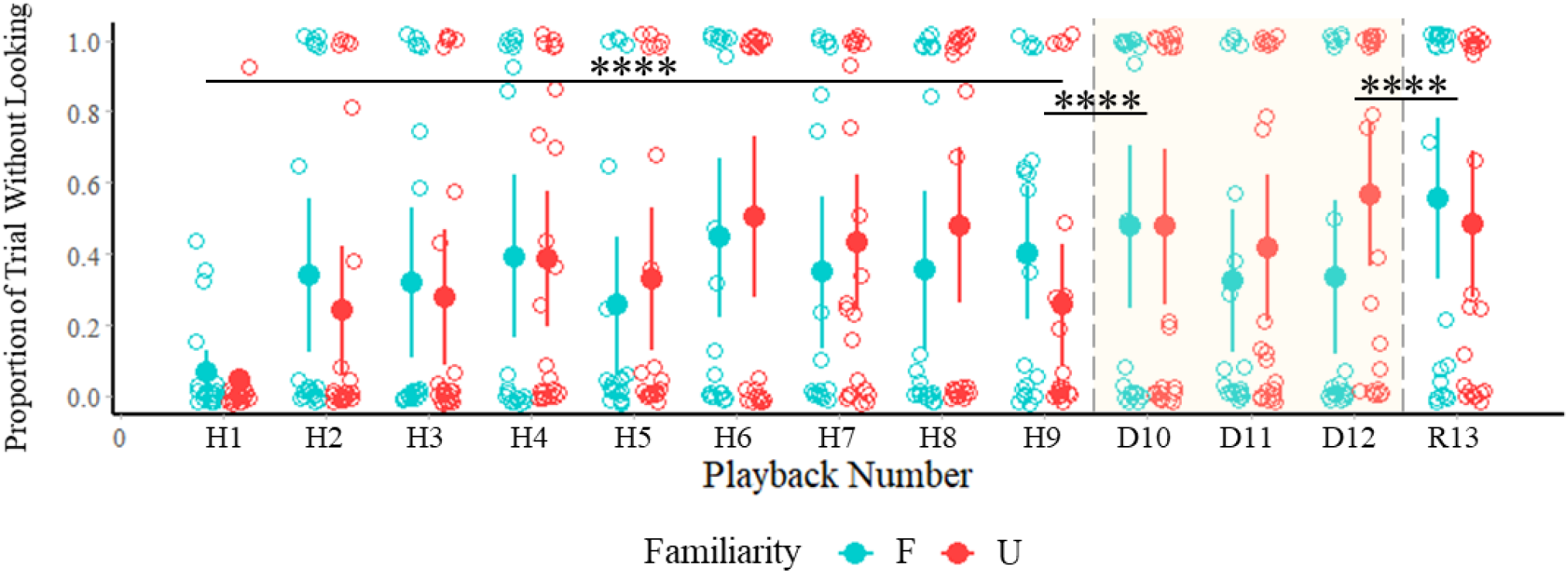
Mean and 95% confidence intervals shown for the effect of playback number and familiarity on goat latency to look following presentation of human voice samples. Playbacks comprising the dishabituation phase (D10-D12) are highlighted and significance of interaction between playback number and familiarity is indicated above. \*\*\*\**p*<0.0001

**Table 2.** Predictors of time goats took to respond after each playback over the habituation phase (H1-H9), following the valence change between the last playback of the habituation and first of the dishabituation phase (H9 v. D10) and to the last playback of the dishabituation phase and to the rehabituation phase (D12 v. R13; binomial GLMM).

| Explanatory Variable | $\beta$ | S.E. | z-value | p-value |
| --- | --- | --- | --- | --- |
| **H1-H9** |  |  |  |  |
| Intercept | -1.884 | 2.115 |  |  |
| **Playback no.** | **0.151** | **0.001** | **294.86** | **<2x10<sup>-16</sup>\*\*\*\*** |
| **Playback no. x Familiarity (U)<sup>a</sup>** | **0.059** | **0.001** | **81.85** | **<2x10<sup>-16</sup>\*\*\*\*** |
| Familiarity (U)<sup>a</sup> | -1.244 | 1.308 | -0.95 | 0.341 |
| Playback Order (HAH)<sup>b</sup> | -0.790 | 1.891 | -0.42 | 0.676 |
| **H9 v. D10** |  |  |  |  |
| Intercept | 3.025 | 1.865 |  |  |
| **Playback no. (D10)<sup>c</sup>** | **0.467** | **0.006** | **82.58** | **<2x10<sup>-16</sup>\*\*\*\*** |
| **Playback no. (D10)<sup>c</sup> x Familiarity (U)<sup>a</sup>** | **1.968** | **0.010** | **195.17** | **<2x10<sup>-16</sup>\*\*\*\*** |
| **Familiarity (U)<sup>a</sup>** | **-4.270** | **1.547** | **-2.76** | **0.006\*\*** |
| Playback Order (HAH)<sup>b</sup> | -4.156 | 2.452 | -1.70 | 0.090 |
| **D12 v. R13** |  |  |  |  |
| Intercept | 0.925 | 2.324 |  |  |
| **Playback no. (R13)<sup>d</sup>** | **2.405** | **0.008** | **283.10** | **<2x10<sup>-16</sup>\*\*\*\*** |
| **Playback no. (R13)<sup>d</sup> x Familiarity (U)<sup>a</sup>** | **-3.322** | **0.011** | **-297.82** | **<2x10<sup>-16</sup>\*\*\*\*** |
| Familiarity (U)<sup>a</sup> | -0.456 | 1.972 | -0.23 | 0.817 |
| Playback Order (HAH)<sup>b</sup> | 2.203 | 3.102 | 0.71 | 0.478 |

Significant results are shown in bold. Key: U = Unfamiliar; HAH = Happy-Angry-Happy Playback Order. Reference Categories: a = Familiar; b = Angry-Happy-Angry playback sequence; c = H9; d = D12. \*\* $p < 0.01$ ; \*\*\* $p < 0.001$ ; \*\*\*\* $p < 0.0001$

Goats were slower to look at the speaker following a change in valence between the habituation and dishabituation phase (H9 v. D10 comparison; playback number x familiarity interaction: β ± s.e. = 1.968 ± 0.010, Z = 195.17, *p*<2x10^-16^****), with how long they took to respond affected by familiarity with the human voice presented (Table 2; Fig. 3). To expand, the increase in time taken to look after D10, compared to H9 was greater when goats were experiencing unfamiliar (25.67% increase; β ± s.e. = -2.435 ± 0.008, z-ratio= -291.68, *p*<0.0001), compared to familiar voices (8.38% increase; β ± s.e. = -0.467± 0.006, z-ratio= - 82.58, *p*<0.0001; see Appendix Table A2 for further *post hoc* comparisons). When considering responses to H9 alone, goats were faster to look when the voice presented was unfamiliar (EMM ± s.e. = 3.48% ± 4.89 into response period), than when it was familiar (EMM ± s.e. = 72.06% ± 29.36; β ± s.e. = 4.270 ± 1.547, z-ratio= 2.76, *p* = 0.030), with no significant differences in time taken to respond to D10 between familiar and unfamiliar voices. Goats were also marginally faster to look when voice valence changed from happy to angry between the habituation and dishabituation phases, than *vice versa* (β ± s.e. = -4.156 ± 2.452, z-value = -1.70; *p* = 0.090).

Goats showed differences in time taken to look between the dishabituation and rehabituation phases, with the direction of this change affected by their familiarity with the voice presented (playback number x familiarity: β ± s.e.= -3.322 ± 0.011, Z = -297.82, *p*<2x10^-16^****; Table 2; Fig. 3). Specifically, goats were slower to look after R13, relative to D12 when the voice was familiar (β ± s.e. = -2.405 ± 0.009, z-ratio= -283.10, *p*<0.0001), but when the voice was unfamiliar, they were slightly faster to respond to R13 (β ± s.e.= 0.916 ± 0.007, z-ratio = 126.82, *p*<0.0001; Appendix Table A2). By contrast, the order in which valence was presented over the habituation, dishabituation and rehabituation phases had no significant effect on how long it took goats to respond after D12, or R13.

### 3.3. Heart Rate and Heart Rate Variability

Overall, goat heart rate showed no significant changes over the course of the habituation phase, but did tend to be lower when there was a longer measurement period available for calculating this response (β ± s.e. = -0.007 ± 0.003, Z = -2.82, *p* = 0.005). In addition, HRV may have decreased with increasing playback number over this phase, although the effect was marginally not significant (β ± s.e. - 0.016 ± 0.008, Z = -1.93, *p* = 0.054; Table 3). We found no further significant changes in heart rate or HRV in relation to changes in valence of voice recordings during the dishabituation phase or when it reversed during the rehabituation phase.

**Table 3.** Predictors of goat heart rate and HRV measured in the 20s response periods following each playback of the habituation phase (H1-H9), following shifts in emotional valence between the habituation and dishabituation phase (H9 v. D10) and the dishabituation and rehabituation phases (D12 v. R13; Poisson GLMM).

| Explanatory Variable | Heart Rate |  |  |  | Heart Rate Variability |  |  |  |
| --- | --- | --- | --- | --- | --- | --- | --- | --- |
| | B | S.E. | z-value | p-value | $\beta$ | S.E. | z-value | p-value |
| **H1-H9** |  |  |  |  |  |  |  |  |
| Intercept | 11.638 | 0.067 |  |  | 10.036 | 0.399 |  |  |
| Playback no. | -0.001 | 0.001 | -0.60 | 0.551 | -0.016 | 0.008 | -1.93 | 0.054 |
| Familiarity (U)<sup>a</sup> | -0.055 | 0.034 | -1.61 | 0.107 | 0.375 | 0.239 | 1.57 | 0.116 |
| Playback Order (HAH)<sup>b</sup> | -0.073 | 0.060 | -1.22 | 0.224 | -0.071 | 0.254 | -0.28 | 0.781 |
| **Measurement Period** | **-0.007** | **0.003** | **-2.82** | **0.005\*\*** | -0.007 | 0.018 | -0.38 | 0.705 |
| **H9 v. D10** |  |  |  |  |  |  |  |  |
| Intercept | 11.645 | 0.131 |  |  | 9.888 | 0.810 |  |  |
| Playback no. (D10)<sup>c</sup> | -0.010 | 0.010 | -1.01 | 0.314 | -0.017 | 0.064 | -0.26 | 0.792 |
| Familiarity (U)<sup>a</sup> | -0.046 | 0.037 | -1.24 | 0.214 | 0.219 | 0.292 | 0.75 | 0.452 |
| Playback Order (HAH)<sup>b</sup> | -0.079 | 0.061 | -1.29 | 0.198 | 0.049 | 0.297 | 0.17 | 0.868 |
| Measurement Period | -0.008 | 0.007 | -1.25 | 0.212 | -0.001 | 0.043 | -0.03 | 0.974 |
| **D12 v. R13** |  |  |  |  |  |  |  |  |
| Intercept | 11.521 | 0.152 |  |  | 11.122 | 1.035 |  |  |
| Playback no. (R13)<sup>d</sup> | -0.001 | 0.010 | -0.06 | 0.956 | 0.003 | 0.082 | 0.04 | 0.970 |
| Familiarity (U)<sup>a</sup> | -0.035 | 0.036 | -0.98 | 0.329 | 0.291 | 0.296 | 0.98 | 0.325 |
| Playback Order (HAH)<sup>b</sup> | -0.067 | 0.059 | -1.12 | 0.261 | -0.020 | 0.284 | -0.07 | 0.944 |
| Measurement Period | -0.003 | 0.008 | -0.42 | 0.677 | -0.073 | 0.055 | -1.34 | 0.182 |

Significant results are shown in bold. Key: U = Unfamiliar; HAH = Happy-Angry-Happy playback order. Reference Categories: a = Familiar; b = Angry-Happy-Angry playback sequence; c = H9; d = D12. \*\* $p < 0.01$

## 4. DISCUSSION

We investigated goat ability to discriminate emotional valence of human speech using a habituation-dishabituation-rehabituation playback paradigm. Having habituated to a human voice conveying a single emotional valence (goats were less likely to look, looked for a shorter time and took longer to look with an increasing number of presentations), goats were less likely to respond following a change in valence (dishabituation phase). However, importantly, those that did look towards the speaker, looked for longer after a change in valence (of the trials in which the subject looked between the habituation and dishabituation phases, in 71.43% goats looked for longer), suggesting these goats had perceived the shift in emotional content of human voice playbacks. After a subsequent reversal of valence to match that of the habituation phase (the rehabituation phase), goats that looked tended to do so for a shorter time than they did after the final playback of the dishabituation phase. Familiarity with the voice presented also seemed to affect goat behavioural responses in complex ways, perhaps indicating a role for familiarity in emotional discrimination, as has been observed in other species (Merola et al., 2014; Briefer et al., 2017). By contrast, changes in goat behaviour with emotional valence, did not seem to be accompanied by shifts in physiological arousal (heart rate and HRV). Goat sensitivity to human emotional cues acts to further emphasize the importance of emotional signalling in regulating social interactions not only between conspecifics (e.g., Schmidt & Cohn, 2001), but also between members of very different species (e.g., Albuquerque et al., 2016; Proops et al., 2018).

Emotions affect vocalisations in similar ways across diverse vertebrate taxa (Briefer, 2020) and resulting similarities in how they are conveyed, in our case between humans and goats, may predispose animals with the ability to discriminate differences in emotional content in vocalisations from other species (Filippi & Gingras, 2018). This is especially the case for emotional arousal, with Filippi et al. (2017) finding human participants could recognize differences in arousal in the vocalisations of nine different species, representing all classes of terrestrial vertebrate. However, goats in our study were required to discriminate human voices based on their valence. Although cross-species evidence for vocal indicators of emotional valence do exist (call duration, fundamental frequency and its variability), these do not appear to apply to the human voice and expression of valence can differ substantially even among closely related species (Briefer, 2012, 2020). Rather than solely relying on broad rules applying across taxa, if the value of information varies according to the heterospecific providing it, we may expect further tuning of social cognitive abilities to interpret the vocal features from more relevant species (e.g., Magrath et al., 2009; Kitchen et al., 2010).

Whether bred for companionship or, like goats, largely for their products, all domesticated animals rely on humans for their habitat, food and protection from predators (MacHugh & Bradley, 2001; Jardat & Lansade, 2021). Through multiple generations of this interdependent relationship, goats and other domesticated species have undoubtedly become well-adapted to occupying this anthropogenic niche, perhaps predisposing them towards more advanced social cognition of our cues (Hare et al., 2005), including emotional ones. This appears to be the case in pigs, with domestic pigs but not wild boar (*Sus scrofa*) behaving differently to human voices based on the valence they express (Maigrot et al., 2022). Given the value of emotional indicators as predictors of behaviour and social motivations, we may expect goats to use these cues to guide their behaviour when interacting with us as they have been seen to do in relation to our attentional cues (Nawroth et al., 2016a). Horses, for example, remember human emotional cues in the long term, with a single presentation of a photograph expressing a positive or negative facial expression being sufficient to cause changes in their behaviour upon encountering the photograph’s subject 3-6 hours later (Proops et al., 2018). Additionally, use of emotional cues in dogs, cats (*Felis catus*) and horses is not limited to guiding behaviour towards the person expressing them, with their responses to an object changing based on whether a familiar person had expressed positive or negative emotions towards it (Merola et al., 2014, 2015; Schrimpf et al., 2020). If goats can discern differences in emotional valence conveyed in human speech, it suggests such cues are important for animals living alongside us, regardless of the specificities of their domestic history (Nawroth et al., 2018; Jardat & Lansade, 2021). These cognitive abilities may not solely be the product of domestic heritage though, with evidence in companion species suggesting emotional recognition can be further developed over an animal’s lifetime (Merola et al., 2014; Briefer et al., 2017).

Goats were overall less likely to respond following a change in emotional valence of human vocal cues suggesting there could be differences among individuals in the ability to perceive this change. Goats at the study site came from a variety of backgrounds, and their individual experiences with human voices of differing valence and their past association with positive or negative events may have affected the salience of, and/or their ability to discriminate between such cues. Indeed, horses and pigs can readily form associations between human vocal cues and positive and negative events, either experienced directly (horses: d’Ingeo et al., 2019) or indirectly, based on maternal experiences during gestation (pigs: Tallet et al., 2016). In dogs, human emotions appear easier to discriminate if they are more relevant to their daily lives (happiness over fear), if expressed by an owner than by a stranger (Merola et al., 2014) and even when shown by individuals of the same gender as their owner (Nagasawa et al., 2011). Similarly, we investigated the effect of familiarity with the speaker and discrimination of their emotional cues.

Goats were slower to respond following a change in valence (dishabituation phase), but importantly this change in looking behaviour was more pronounced in those experiencing unfamiliar voices. The relative stability in time taken to respond could suggest attention was retained to a greater extent in goats listening to familiar voices. Subjects also tended to spend less time looking following a reversal of valence (rehabituation phase), suggesting a drop in behavioural arousal. According to Charlton et al. (2007, 2011), this would suggest that the observed increase in looking duration at the start of the dishabituation phase was robust and not due to a chance renewal in attention. Although this decrease in time spent looking after the rehabituation phase was only marginal, taken together with changes in latency to respond, it could indicate that familiarity affects the ease with which goats can discriminate human emotional cues. Specifically, goats experiencing familiar voices were slower to respond following the rehabituation phase, which is consistent with the expected decline in behavioural arousal. Comparatively, goats were actually quicker to respond to unfamiliar voices after a reversal of emotional valence. In sum, as time taken to respond was more stable before and after a change in valence and was longer following a subsequent reversal of valence in goats listening to familiar voices, this could suggest they were better able to discriminate emotional valence in familiar compared to unfamiliar human voices.

Goats took an increasingly longer time to respond as the habituation phase progressed, with this rate of increase being greater when the voice presented was unfamiliar, which could indicate familiar voices were harder to habituate to. However, voice familiarity only weakly affected goat responses over this phase, and when the voice was unfamiliar, goats looked for longer after the end of the habituation and start of the dishabituation phases and were quicker to respond to the former than when the voice was familiar. Dogs and cats have been shown to monitor emotional displays produced by their owner for longer than those of a stranger, with this difference speculated to have an effect on the relative ease with which animals could interpret their owner’s emotional cues (Merola et al., 2014; Galvan & Vonk, 2016). However, goats in the current investigation did not show robust evidence for being more attentive to familiar over unfamiliar voices. Alternatively, familiarity with a person and, by extension, how they express emotions may facilitate discrimination of these cues, enabling goats to distinguish even more subtle emotional signals (Preston & De Waal, 2002; Briefer et al., 2017). Ultimately though, goats took longer to respond following a change in valence both when the voice was familiar and unfamiliar, which goes against the predicted renewal of attention should they have perceived this change (Charlton et al., 2007, 2011). Familiarity may facilitate the ability to discriminate valence of human voices, but for the current investigation it appears familiarity affects goat behaviour in complex ways which are difficult to interpret.

Further explanations as to why some goats responded for longer and some not at all following a change in voice valence may include variation in cognitive abilities among goats to perceive human emotional cues. Goats may have also needed to maintain a certain level of attention to the playback sequence to discriminate its emotional content. Alternatively, variation in goat behavioural responses could at least in part be an artefact of the playback composition itself. Emotional arousal, as well as the clarity and intensity in which valence was portrayed will have varied among the eight speakers, and between voice samples taken from the same speaker. Although a valence validation test was performed in human participants who were largely able to correctly score emotional valence in voice samples used for testing goats, the ability of these participants to detect differences in valence would have been far more refined than that of a goat. It is therefore possible that valence portrayed in the voice samples was not always clear enough for goats to effectively discriminate.

Our voice can act as a potent and long-range signal of human activity, and has been shown to have sweeping implications for behaviour and habitat use in animal populations living in human-dominated landscapes (McComb et al., 2014; Clinchy et al., 2016; Suraci et al., 2019). The impacts of the human voice on animal behaviour and emotional experiences are no more so true than in the domestic environment where it is heard during daily management practices (e.g., feeding and cleaning), although its implications for animal welfare remain poorly understood (Tallet et al., 2018). Voices conveying negative valence (e.g., shouting, use of a stern tone and growling) have been shown to produce fear and vigilance-related responses in domesticated animals (Merkies et al., 2013; Smith et al., 2018), whereas soothing voices may be calming (Lange et al., 2020). Furthermore, positive human emotional cues have been linked to approach, including in goats (Nawroth et al., 2018), so may facilitate human-animal bonding. Although goats in the current investigation did not show clear changes in physiological arousal linked to emotional valence of human voices, we only looked at cardiac responses and did not see how these changed with valence over playback phases (e.g., habituation phase). It is therefore still possible an emotional response took place, which may have been detectable using a different behavioural or physiological measure. Further research is needed to understand the importance of the human voice on the emotional lives and welfare of goats and other domesticated species.

In summary, we present here the first evidence that goats can discriminate between cues expressed in the human voice, namely, emotional valence. These findings contribute to the limited literature available indicating livestock, like companion animals, are sensitive to human emotional cues (e.g., Nawroth et al., 2018; Destrez et al., 2021; Maigrot et al., 2022). Our findings raise a number of questions, not limited to those regarding the role of domestication on the development of these abilities. Indeed, the observed differences in goat responses to human emotional cues may stress the importance of individual experiences and learning in particular on interspecific emotional communication (Nagasawa et al., 2011; Merola et al., 2014; Tallet et al., 2016; d’Ingeo et al., 2019).

## AUTHOR CONTRIBUTIONS

**Marianne A. Mason:** Conceptualisation, Data curation, Formal analysis, Investigation, Methodology, Project administration, Visualisation, Original draft preparation. **Stuart Semple:** Conceptualisation, Funding acquisition, Methodology, Supervision, Writing-Review & Editing. **Harry H. Marshall:** Formal analysis, Supervision, Visualisation, Writing-Review & Editing. **Alan G. McElligott:** Conceptualisation, Funding acquisition, Methodology, Supervision, Writing-Review & Editing.

## ACKNOWLEDGEMENTS

We are very grateful to all our field assistants and to Anastasia Makhlouf, Tania Perroux, Dr Luigi Baciadonna and Dr Elodie Briefer especially, and to everyone who took part in our valence validation experiment. We also thank everyone at Buttercups Sanctuary for Goats for their advice and for allowing us to access the goats.

---

**Figure A1.**
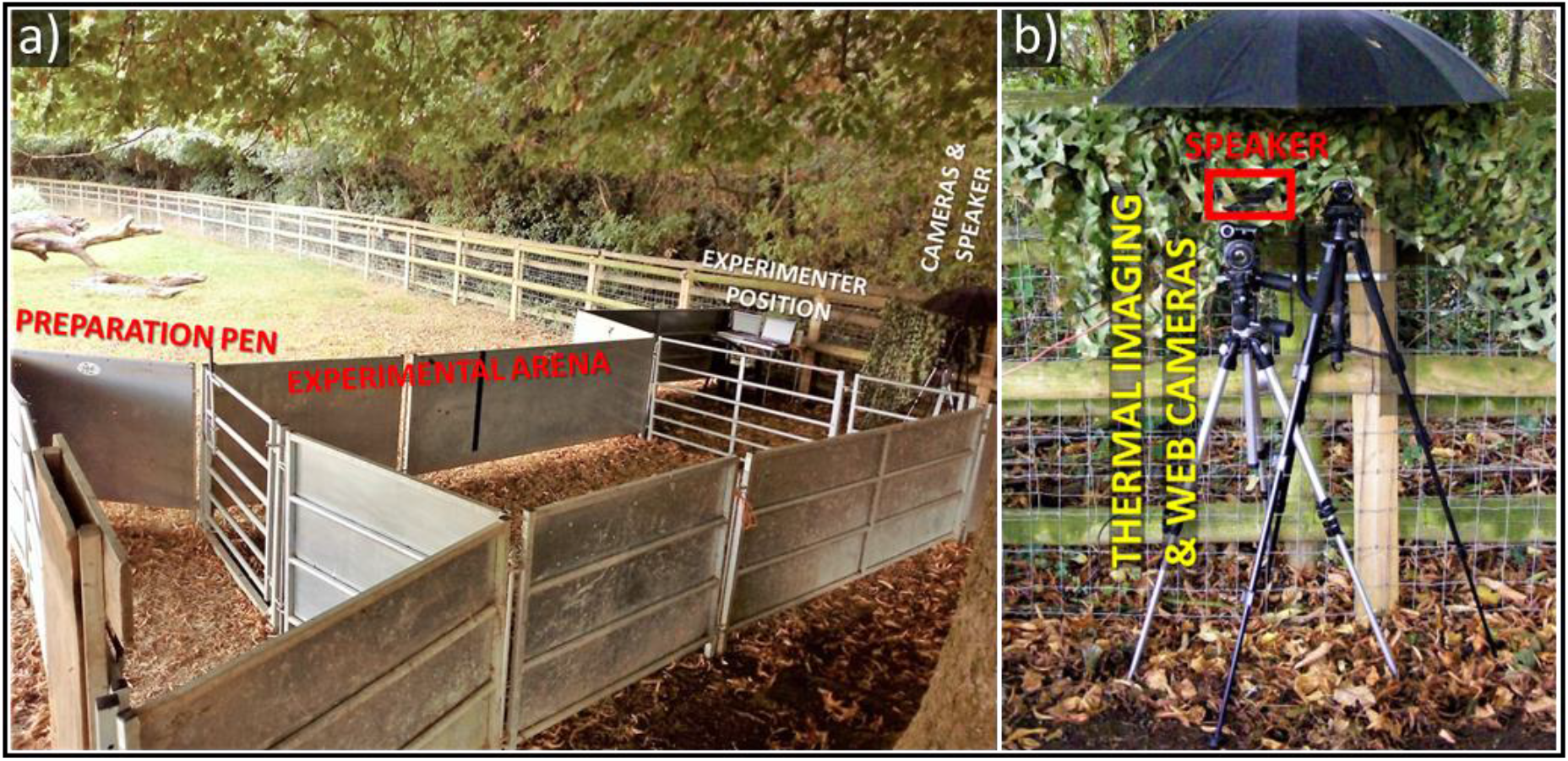
a) View of experimental enclosure indicating the relative locations of the preparation pen, where goats were fitted with a heart rate monitor, the experimental arena, where goats were held during experimental trials, as well where the experimenter, speakers and video camera would have been positioned. b) Set-up of the speaker, video camera and back up web camera. We initially planned to collect thermal imaging data, however, attempts to get thermal images of sufficient quantity and quality were ultimately unsuccessful.

**Table A1.** Subject name, sex (F= Female; M= Male), breed, age, number of years at the study site and playback sequence experienced during experimental trials (AHA = Angry-Happy-Angry; HAH = Happy-Angry-Happy)

| Goat ID | Sex | Breed | Age (years) | Time at Sanctuary (years) | Playback Sequence |
| --- | --- | --- | --- | --- | --- |
| Anoushka | F | Pygmy | 14+ | 2 | AHA |
| Arlo | M | Mixed Breed | 7 | 3 | HAH |
| Bear | M | Pygmy | 6 | 2 | AHA |
| Bernard | M | Anglo Nubian | 8 | 8 | AHA |
| Bramble | M | Pygmy | Unknown | 1 | HAH |
| Bonnie | F | Pygmy | 16 | 6 | AHA |
| Buttercup | F | Pygmy | ≈14 | 1 | HAH |
| Davey | M | Saanen | 14 | 6 | AHA |
| Del Boy | M | Pygmy | 10 | 6 | AHA |
| Dixie | F | Pygmy | 10 | 8 | HAH |
| Ewok | M | Anglo Nubian | 3 | 2 | HAH |
| Harry | M | Saanen | 10 | 8 | HAH |
| Heidi | F | Toggenburg | 9 | 7 | HAH |
| Luke | M | Anglo Nubian | 3 | 2 | HAH |
| Natalie | M | Mixed Breed | 9 | 8 | HAH |
| Pickle | F | Pygmy | Unknown | 2 | AHA |
| Princess | F | Anglo Nubian | 3 | 2 | HAH |
| Rambo | M | Unknown | 7 | 2 | HAH |
| Rosie | F | Saanen | 7 | 1 | HAH |
| Sapphire | F | Toggenburg | 8 | 3 | AHA |
| Shakespeare | M | Saanen – Golden Guernsey Cross | 12 | 9 | HAH |
| Sticky | M | Saanen – Golden Guernsey Cross | 6 | 6 | AHA |
| Sugar | F | Anglo Nubian | 7 | 2 | AHA |
| Tarnie | F | Golden Guernsey | 15 | 3 | AHA |
| Valentine | F | Toggenburg | Unknown | 2 | AHA |
| Vern | M | Toggenburg | 8 | 8 | AHA |
| Wendy | F | Toggenburg | Unknown | 1 | HAH |

**Table A2.**
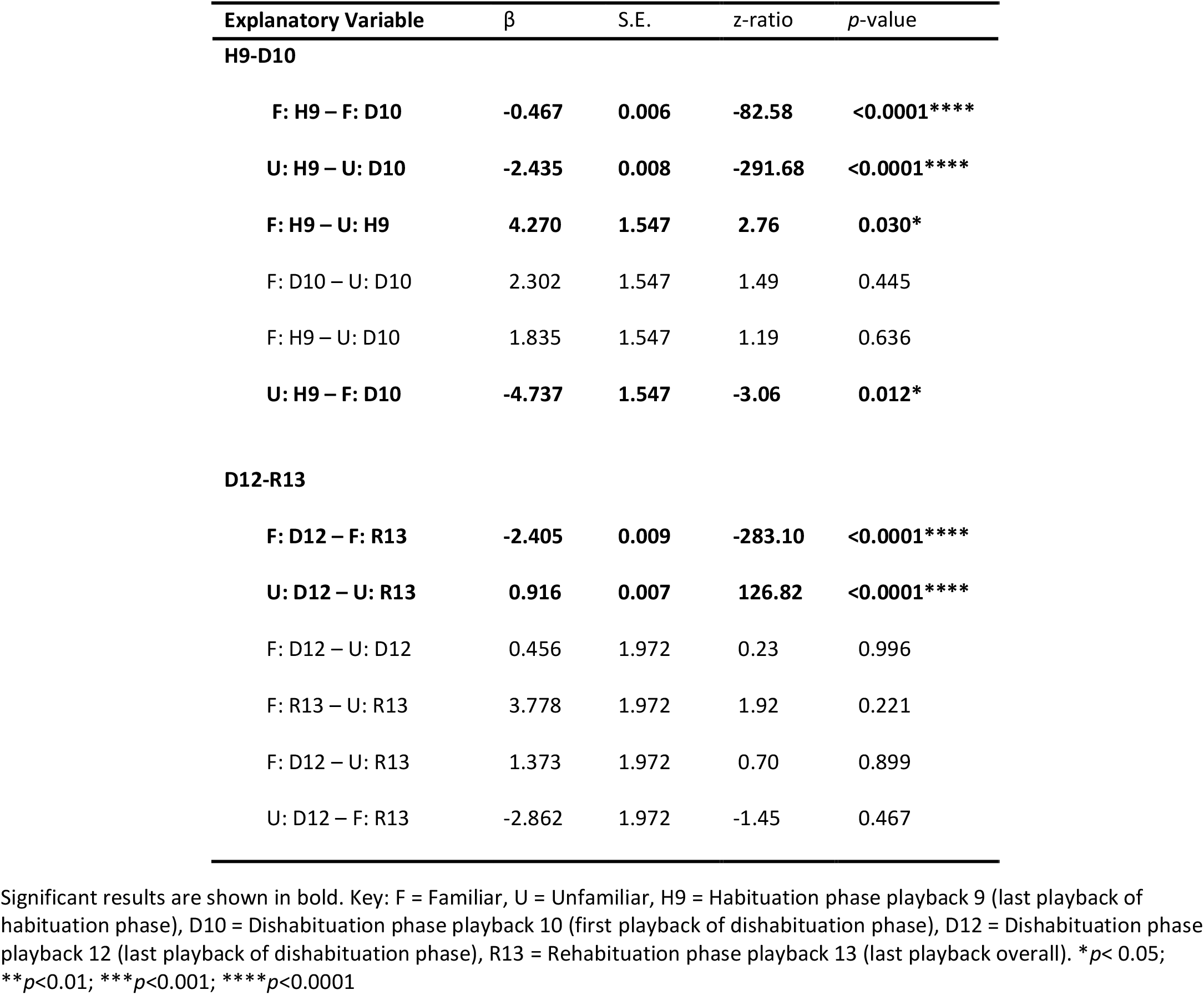
Results of *post hoc* comparisons for the interaction between playback number and familiarity on goat latency to look at the sound source following a change in emotional valence of human voice samples between the habituation and dishabituation phase (H9-D10) and the dishabituation and rehabituation phases (D12-R13).

| Explanatory Variable | $\beta$ | S.E. | z-ratio | p-value |
| --- | --- | --- | --- | --- |
| **H9-D10** |  |  |  |  |
| **F: H9 – F: D10** | **-0.467** | **0.006** | **-82.58** | **<0.0001\*\*\*\*** |
| **U: H9 – U: D10** | **-2.435** | **0.008** | **-291.68** | **<0.0001\*\*\*\*** |
| **F: H9 – U: H9** | **4.270** | **1.547** | **2.76** | **0.030\*** |
| F: D10 – U: D10 | 2.302 | 1.547 | 1.49 | 0.445 |
| F: H9 – U: D10 | 1.835 | 1.547 | 1.19 | 0.636 |
| **U: H9 – F: D10** | **-4.737** | **1.547** | **-3.06** | **0.012\*** |
| **D12-R13** |  |  |  |  |
| **F: D12 – F: R13** | **-2.405** | **0.009** | **-283.10** | **<0.0001\*\*\*\*** |
| **U: D12 – U: R13** | **0.916** | **0.007** | **126.82** | **<0.0001\*\*\*\*** |
| F: D12 – U: D12 | 0.456 | 1.972 | 0.23 | 0.996 |
| F: R13 – U: R13 | 3.778 | 1.972 | 1.92 | 0.221 |
| F: D12 – U: R13 | 1.373 | 1.972 | 0.70 | 0.899 |
| U: D12 – F: R13 | -2.862 | 1.972 | -1.45 | 0.467 |

Significant results are shown in bold. Key: F = Familiar, U = Unfamiliar, H9 = Habituation phase playback 9 (last playback of habituation phase), D10 = Dishabituation phase playback 10 (first playback of dishabituation phase), D12 = Dishabituation phase playback 12 (last playback of dishabituation phase), R13 = Rehabilitation phase playback 13 (last playback overall). \* $p < 0.05$ ; \*\* $p < 0.01$ ; \*\*\* $p < 0.001$ ; \*\*\*\* $p < 0.0001$

